# MiCoP: Microbial Community Profiling method for detecting viral and fungal organisms in metagenomic samples

**DOI:** 10.1101/243188

**Authors:** Nathan LaPierre, Serghei Mangul, Mohammed Alser, Igor Mandric, Nicholas C. Wu, David Koslicki, Eleazar Eskin

## Abstract

**Background:** High throughput sequencing has spurred the development of metagenomics, which involves the direct analysis of microbial communities in various environments such as soil, ocean water, and the human body. Many existing methods based on marker genes or k-mers have limited sensitivity or are too computationally demanding for many users. Additionally, most work in metagenomics has focused on bacteria and archaea, neglecting to study other key microbes such as viruses and eukaryotes.

**Results:** Here we present a method, MiCoP (Microbiome Community Profiling), that uses fast-mapping of reads to build a comprehensive reference database of full genomes from viruses and eukaryotes to achieve maximum read usage and enable the analysis of the virome and eukaryome in each sample. We demonstrate that mapping of metagenomic reads is feasible for the smaller viral and eukaryotic reference databases. We show that our method is accurate on simulated and mock community data and identifies many more viral and fungal species than previously-reported results on real data from the Human Microbiome Project.

**Conclusions:** MiCoP is a mapping-based method that proves more effective than existing methods at abundance profiling of viruses and eukaryotes in metagenomic samples. MiCoP can be used to detect the full diversity of these communities. The code, data, and documentation is publicly available on GitHub at: https://github.com/smangul1/MiCoP

## Background

Microorganisms are ubiquitous in almost every ecosystem on earth, including soil, ocean water, and the human body. Single-celled organisms play a number of vital roles in each of these environments [1,2]. Identifying the microbes present in a sample is critical to understanding what functions are carried out by these organisms and characterizing how disturbances in microbial communities lead to various maladies. Traditionally, microorganisms have been studied via culture-based techniques, in which the microbial organisms were isolated and studied individually in laboratory settings. However, it is well-recognized that many microbes are not culturable; hence they cannot be studied in laboratory settings [3]. In addition, techniques studying microbes in laboratory settings are incapable of capturing the complex relations between hundreds to thousands of different microbial species in their natural habitats [1,2]. High-throughput sequencing has revolutionized microbiome research, enabling the study of thousands of microbial genomes directly from their host environments and forming the field of metagenomics. This approach bypasses the traditional culture-dependent bias and allows the study of the composition of microbial communities in their natural habitats across different human tissues and environmental settings [1, 2]. Metagenomic profiling has proven useful for studying various microbes, including eukaryotic and viral pathogens, which were previously impossible to study in an unbiased manner with 16S ribosomal RNA gene sequencing [4–6].

Despite the critical importance of the “virome” and the “eukaryome” in affecting the microbiome and human health, most metagenomic profiling methods have focused primarily on identifying bacteria and archaea [7,8]. Several existing methods for metagenomic profiling have proposed using ‘marker genes’ that uniquely identify a read as coming from a certain species. This method has been shown to be efficient and accurate at estimating the presence and relative abundances of bacteria and archaea in a sample [9–11]. However, approaches based on marker genes have some limitations with identifying viral and eukaryotic genomes. One approach involves comparing differences in genes that are considered ‘universal’ but differ between species. This approach uses reads that indicate a certain sequence for that marker gene and thus uniquely identify a species [9]. However, this is problematic for viruses, which are comprised mostly of novel sequences and do not share any single common gene [2,12,13]. Another approach utilizes sequences that uniquely identify a given clade [10,11], but these can only use the relatively small number of reads that map to these specific regions of the genome [14], leading to poor sensitivity [15]. This is particularly problematic for eukaryotic genomes, which are usually long and comprised mostly of noncoding regions, leading to poor read utilization [5,16,17]. Recent approaches based on k-mers have overcome these issues and improved run time dramatically [14,18]. However, these approaches show decreased sensitivity due to requiring perfect k-mer matches [15,19]. In addition, they demand heavy memory usage often in excess of 100GB, which many users do not have available [20]. Finally, k-mer based methods have been observed to predict a large number of low-abundance species that are not actually present in the sample [15].

In this paper, we present MiCoP (MIcrobiome COmmunity Profiling), a computational method capable of profiling viruses and eukaryotes with high precision and sensitivity. We overcome the issues mentioned above by utilizing a fast mapping-based approach, which is capable of high read usage, avoids bias against viruses and eukaryotes, and is sensitive to low-abundance species. Mapping-based approaches have been observed to have even higher sensitivity than Megablast [21], a common gold standard method, but there have traditionally been concerns with the speed and memory usage of a mapping-based approach [22]. We demonstrate that, when using smaller viral and eukaryote reference databases, a mapping-based approach is both feasible and preferable.

We first map reads to reference genomes using BWA, and we then apply a two-step read assignment process. From the BWA mapping results, uniquely-mapped reads are immediately assigned, and, subsequently, multi-mapped reads are probabilistically assigned to genomes based on the distribution of uniquely-mapped reads. We perform species abundance estimation by calculating the number of reads mapped to each genome and normalizing by genome length. Since MiCoP maps reads to specific genomes in a reference database, it is capable of detecting microbes at a finer granularity than the species level, for instance if different strains or chromosomes from a species are listed separately in the reference database. We validate MiCoP by comparing its abundance estimation performance with two of the most popular methods, MetaPhlAn2 [11] and Kraken [14]. We demonstrate improved results on simulated reads from viral and eukaryotic genomes, and we show that MiCoP can identify more viruses and eukaryotes in Human Microbiome Project data than previously-used methods.

## Results

### Methods overview

MiCoP utilizes a mapping-based approach to perform highly sensitive and precise read classification and accurate abundance estimation of viruses and eukaryotes in metagenomic samples. MiCoP starts with mapping reads to whole genomes in a reference database using BWA [23], keeping all multi-mapped reads. Our approach then use a two-stage process to classify the reads. In the first stage, all uniquely-mapped reads are classified, and we compute the abundance of each genome in the sample based on these reads. In the second stage, multi-mapped reads are probabilistically assigned to one of the genomes that they mapped to, with probabilities proportional to the abundance of those genomes among uniquely-mapped reads. We remove species for which there is limited evidence, based on the number of reads assigned to that species. Relative abundances of the present genomes are then computed. These steps are discussed in further detail in the methods section. Figure 1 illustrates the MiCoP workflow.

**Fig. 1.**
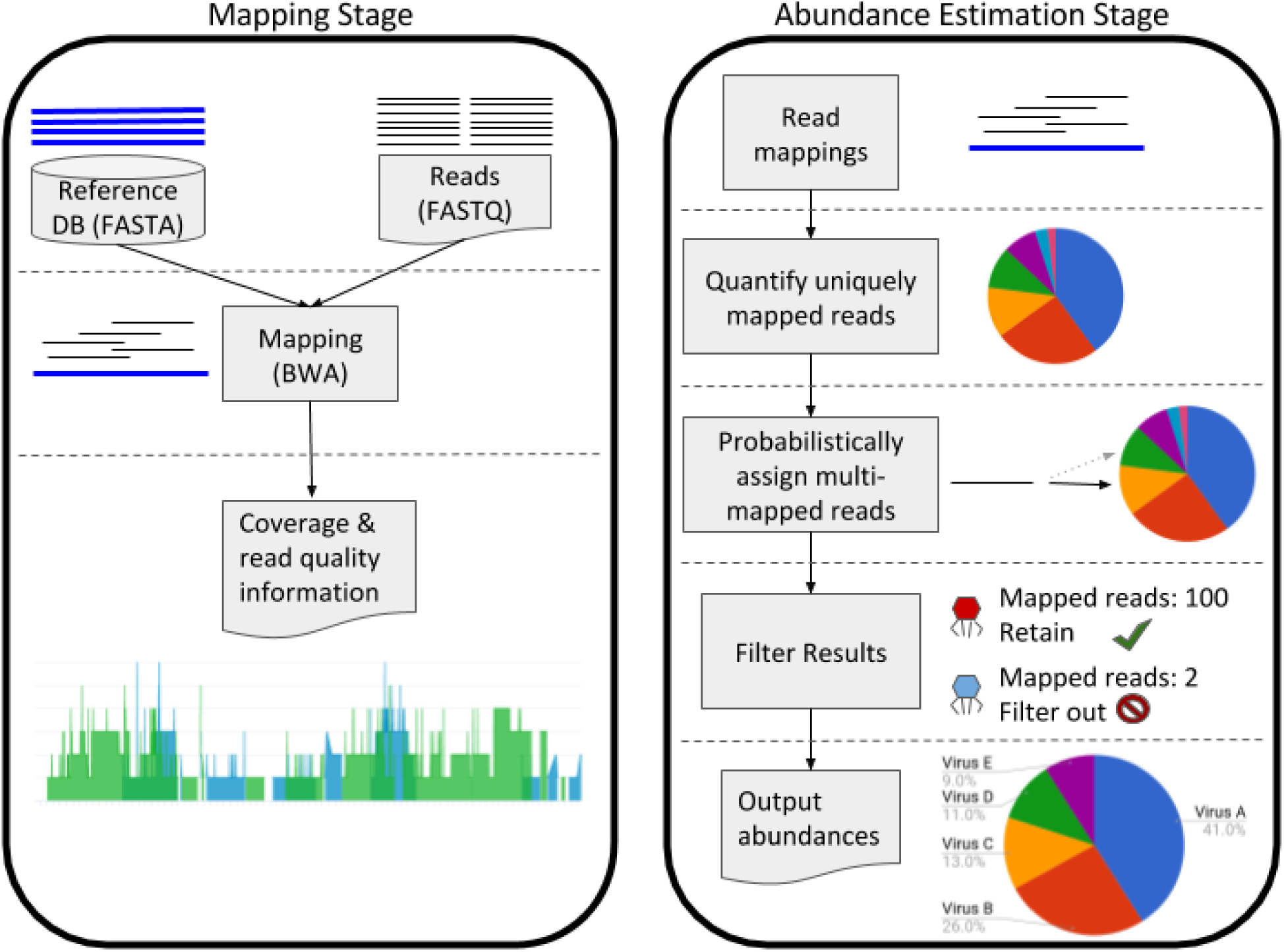
MiCoP workflow. Reads are first aligned to viral or eukaryotic genomes in a reference database using BWA. The results provide coverage and read mapping quality information that can be examined. In the abundance estimation stage, uniquely-mapped reads are assigned to species and species abundances are estimated based on these. Multimapped reads are then assigned to genomes with probability proportional to their abundances among uniquely-mapped reads. Species with not enough reads mapped are filtered out, and then the final species abundances are computed.

### Performance Metrics

We evaluate the performance of different methods using several different metrics, which are designed to encompass both performance in the binary classification task of predicting species presence or absence and in the estimation of relative abundances. For species presence and absence, a “True Positive” (TP) indicates that a species that is actually present in a sample is correctly predicted as being present by a method, while a “False Positive” (FP) indicates that the method predicted the presence of a species that is not actually in a sample and a “False Negative” (FN) indicates that a species was actually present in a sample but a method did not predict its presence. We use two metrics to assess the performance of a method in species presence/absence, precision and recall, defined below. Precision measures the percentage of predicted species that are actually present, while recall measures the percentage of species actually in a sample that were predicted by a method. Additionally, we report the F1-Score, which is defined as the harmonic mean of precision and recall. All three of these metrics range from 0 to 1, or 0% to 100%.

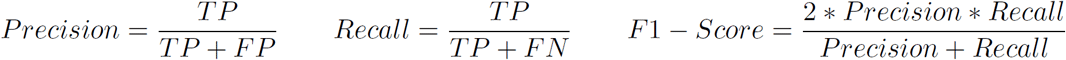

We use the L1 Error as a measure of how accurately a method computes the relative abundances of species in a sample. The L1 Error is the sum of absolute value differences between predicted species abundances and actual species abundances, and ranges from 0 (completely correct) to 2 (completely incorrect). An L1 Error of 0 indicates that the exact set of actual species and their actual abundances is predicted perfectly, while a score of 2 indicates that the set of predicted species is completely incorrect. The L1 Error can be described mathematically as:

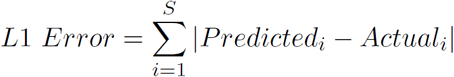

where S is the set of species that are predicted or actually present and i is the summation index.

### MiCoP shows order of magnitude improvement in abundance estimation

We validated the accuracy of our method by using simulated data, so that our results could be compared to a known ground truth. We sampled 1 million reads from 544 viral genomes obtained from an NCBI RefSeq reference file database 5808 viral genomes. Our simulation was designed using the “high complexity” microbial community parameters specified by the CAMI consortium, as described in the methods section [15]. While 1 million reads is a fairly small metagenomic sample, in our case the coverage was reasonable because viral genomes are much shorter than bacterial genomes. We compared the results of our method with two of the most popular metagenome profiling methods, MetaPhlAn2 [11] and Kraken [14]. For this initial simulation, we used the default MetaPhlAn database and Kraken’s pre-built Minikraken database, since these databases reflect the most common conditions under which these methods are applied. For MiCoP, we use a database composed of the genomes available from NCBI’s RefSeq Viral and Fungal databases. In our next simulation, we examine the effect of the choice of reference database. Results are shown in Table 1.

**Table 1.**
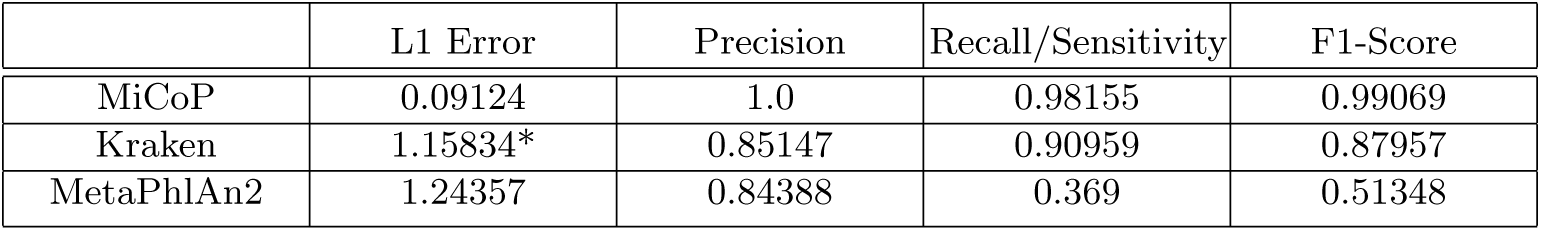
Abundance estimation performance results on a simulated viral community with 544 species. Kraken is considered by its authors to be a read classification tool, not abundance estimation tool, so we put an asterisk next to its results. However, we note that abundance estimation is a common application for Kraken in practice. Overall, MiCoP outperforms the other two methods across all metrics. Kraken and especially MetaPhlAn are limited by the poor representation of viruses in their standard databases. L1 error is the sum of the absolute values of the differences between the computed species abundances and the ground truth species abundances. MiCoP’s L1 error was more than an order of magnitude better than the other tools, and MiCoP had the best precision and recall.

We found that all three methods had high precision, but MiCoP had perfect precision. In other words, every species MiCoP predicted as present in the sample was actually present according to the ground truth. Results from MetaPhlAn2 reported only 37% sensitivity, indicating that it only identified just over a third of the species present in the sample, while Kraken identified about 91% and MiCoP identified about 98%. The total L1 error was only about 0.09 for MiCoP, while MetaPhlAn2’s error was over 1.24. Kraken also reported a high L1 error, but its authors present Kraken as a read classification method, not a relative abundance estimation method, so this metric may be misleading for Kraken.

Clearly, MetaPhlAn2’s performance was limited by the fact that their standard database did not contain marker genes for many of the NCBI virus genomes present in the sample. When using MetaPhlAn2’s provided database, researchers may fail to identify many of the species present in a sample, simply because they are not in this database. While Kraken’s default Minikraken database seems more comprehensive, it is still known to have lower sensitivity when compared to a more complete reference [14]. The problem of reference bias can significantly impact the performance of these methods, particularly when applied to real datasets in which the set of expected genomes is not known in advance.

However, for the purposes of comparing MiCoP to MetaPhlAn2 and Kraken without the results being affected by reference bias, we constructed a dataset composed only of genomes that all three of these methods identified in the high complexity dataset. Out of these 173 genomes, we selected 40 according to the “low complexity” microbial community parameters established by the CAMI consortium [15]. We also simulated errors in these reads, with the error rate linearly increasing from 1% at the start of reads to 5% at the end, with 2/3 of errors being substitutions and the other 1/3 being indels; these numbers were chosen to be roughly equal to the error rates used in the Kraken paper [14]. Results are shown in Table 2.

**Table 2.** Abundance estimation performance results on a simulated viral community with 40 species. These species were sampled from the intersect of the species detected by all three tools in the previous simulation. Thus, this simulation consisted of only the species that were present in all three reference databases, eliminating reference bias. MetaPhlAn’s performance dramatically improved, predicting the exact set of species in the sample, but its abundance estimation was an order of magnitude worse than MiCoP’s. Kraken’s results did not markedly improve in this simulation.

|  | L1 Error | Precision | Recall/Sensitivity | F1-Score |
| --- | --- | --- | --- | --- |
| MiCoP | 0.00909 | 1.0 | 1.0 | 1.0 |
| Kraken | 1.15466* | 0.82222 | 0.925 | 0.87059 |
| MetaPhlAn2 | 0.09844 | 1.0 | 1.0 | 1.0 |

We observed that MiCoP and MetaPhlAn2 both identified the exact set of genomes present in the sample, leading to perfect precision and recall scores. However, MiCoP’s L1 error of about 0.0091 was less than a tenth of MetaPhlAn2’s L1 error of about 0.098. We speculate that the poor read utilization of MetaPhlAn2 (about 5% of reads used) leads to less accurate abundance estimation. Kraken’s sensitivity was almost perfect, with only three genomes out of 40 left unidentified. Surprisingly, Kraken’s precision was surprisingly slightly worse in comparison to its performance on the high complexity dataset. We observed that Kraken reported many low-abundance false positive predictions, resulting from no more than a few mispredicted reads. This phenomenon was also reported by the CAMI consortium when testing on bacterial data [15]. By excluding genomes that were reported in less than 0.01% of reads by Kraken, we were able to raise precision from the original 0.39362 to 0.82222 without reducing recall. With higher cutoffs, the recall dropped rapidly, so the choice of 0.01% appeared to be optimal. Even with this improved precision, Kraken still produces several false positives whereas the other methods do not. These results suggest that MiCoP is highly effective at accurately predicting the species present in a sample and estimating their relative abundances, even when a significant amount of errors are present in the reads.

We generated a low complexity fungi community simulation dataset using the procedure described above: first we simulated a high complexity dataset, and then sampled 40 genomes out of the genomes detected by all methods on the high complexity dataset. Unlike the viral simulation, in which each sampled genome belongs to a different viral species, this simulation’s 40 genomes derived from only 7 different fungal species (some genomes in the reference database were contigs from the same species). Results are shown in Table 3.

**Table 3.**
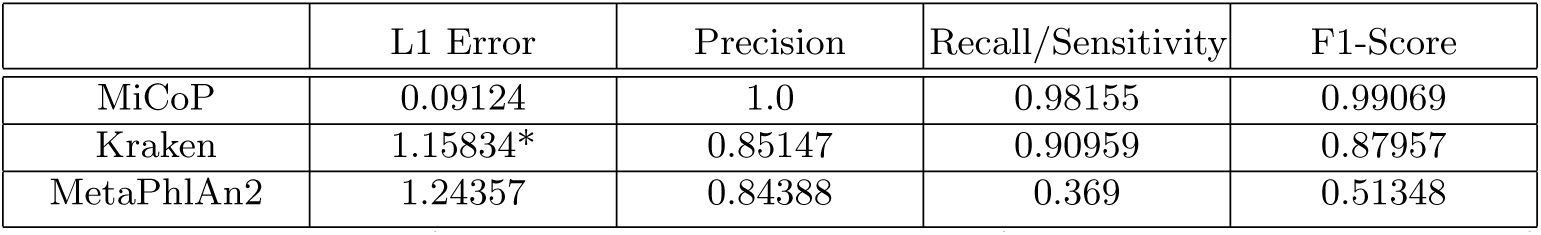
Abundance estimation performance results on a simulated fungal community consisting of 40 genomes derived from 7 different species. These species were sampled in the same way as in the previous table: by taking the intersect of species detected by all three tools on a higher-complexity community. MiCoP detected the exact set of species present in the sample, while Kraken had one false negative and MetaPhlAn had one false positive. Additionally, MiCoP’s abundance estimation was more than an order of magnitude better than the other tools.

Due to the relatively small number of species present, with several genomes sampled from each species, all methods were able to predict almost the exact set of species present. MetaPhlAn generated one false positive and Kraken generated one false negative. However, MiCoP achieved an L1 error that was more than 10 times lower than that of Kraken or MetaPhlAn2. This result demonstrates that MiCoP is effective at estimating the relative abundance of eukaryotes in a sample more accurately than existing methods, even when those methods predict almost the exact set of species in the sample correctly.

### MiCoP detects greater diversity of viruses and eukaryotes in real world data

The Human Microbiome Project (HMP) is an ongoing large-scale effort to understand and characterize the human microbiome across a variety of body sites [24–26]. One of the main HMP studies took 4788 samples from 300 patients across 18 body sites [26]. Notably, this study used MetaPhlAn to profile their metagenomic samples [26]. As our simulations indicate, it is possible that analyses using MetaPhlAn have failed to capture the diversity and prevalence of the human virome and eukaryome due to a functional bias towards bacteria. We have previously shown MiCoP’s superior performance to MetaPhlAn on simulated datasets. However, real data may not be as clean as simulated data, due to factors such as library preparation, mutations in organisms in real world environments, horizontal gene transfer, etcetera. We have shown that Kraken leads to a number of false positives even in the simulated data, and thus we do not assess its results on real data where ground truth is not available.

In order to compare the performance of MiCoP against MetaPhlAn2 on real world data, we applied both methods to publicly-available mock community data. Mock communities have the advantage of being examples of real live microbiomes, but with the community composition controlled and known in advance, providing an effective means for evaluating performance on real world data. Additionally, because classic mapping methods such as BWA were not originally designed for metagenomics [22], they can occasionally assign reads incorrectly when applied to noisier real world data, especially when organisms in the sample are not in the reference database but closely related organisms are. Mock communities thus provide an opportunity to establish parameter settings for species filtering that perform well on real data. Several large scale and high profile metagenomics studies have applied BLAST or other alignment methods with a variety of different parameter settings with little to no explanation [27–29], so establishing a standard for these settings for our method is useful and illustrative. We focus on the precision and recall metrics for the mock community comparisons, as the studies that the communities were taken from report the abundances of the species in ways that could not be reliably translated into normalized relative abundances. As with our simulation studies, we use a database composed of the genomes available from NCBI’s RefSeq Viral and Fungal databases. We focus on fungi in particular because RefSeq’s databases for non-fungal eukaryotes are currently very limited. For MetaPhlAn2, we use their provided marker gene reference database.

We first applied MiCoP and MetaPhlAn2 to a viral mock community consisting of 9 species that was released by Conceio-Neto *et al.* in 2015 [30]. We empirically found that the optimal parameter settings for MiCoP required at least 10 reads with 60% of bases mapped to the reference genome to consider a species present. Results are shown in Table 4. MetaPhlAn only detected 1 of the 9 species in the sample, with no false positives, while MiCoP detected 7 of the 9 species in the sample with 1 false positive. The one false positive that MiCoP detected occurred due to the previously-mentioned reference bias problem. The species that was actually present in the sample, Feline panleukopenia virus, was not in the NCBI viral reference database, while the closely-related [31] Canine parvovirus was, leading to many of the reads from Feline panleukopenia virus mapping well to the Canine parvovirus genome. While very stringent parameter settings could filter out this false positive, they would also filter out many truly present species from the results. This result highlights both the reference bias problem and the inherent tradeoff between false positive and false negative rates. Regardless, MiCoP represented the viral community much more accurately than MetaPhlAn2, which failed to capture the community diversity. Finally, in terms of speed, MetaPhlAn2 processed the reads about twice as quickly as MiCoP, although MiCoP still processed the 12.4 million reads in 2 hours and 22 minutes.

**Table 4.** Comparison of the performance of MiCoP and MetaPhlAn2 on a mock viral community consisting of 9 species. MetaPhlAn2 only detects 1 of 9 species, with no false positives, while MiCoP detects 7 of 9 species with one false positive, thus profiling the community much more accurately. MetaPhlAn2 processed the reads about twice as fast as MiCoP.

|  | Precision | Recall | F1-Score | Reads per minute |
| --- | --- | --- | --- | --- |
| MiCoP | 0.875 | 0.77778 | 0.82353 | 87629 |
| MetaPhlAn2 | 1.0 | 0.11111 | 0.19999 | 162845 |

We also applied MiCoP and MetaPhlAn2 to a fungal mock community consisting of 20 species from 4 genera that was released by Tonge et *al.* in 2014 [32]. NCBI’s fungal RefSeq database contains far fewer species than its viral counterpart, and was missing many of the species in the mock community. Thus, species-level results identified several false but closely-related species in the sample, similarly to the previously explored Canine parvovirus example, and we determined that classification of real fungal communities can generally only be done accurately at the genus level. We applied somewhat stricter parameter settings than were applied for viruses, owing to the greater amount of shared genomic sequence between different fungi as compared with viruses. In particular, we considered a genus present if at least 100 reads matched at least 99% with any of the reference genomes for that genus with a maximum of 1 indel or substitution per read. Results are shown in Table 5. MetaPhlAn2 did not detect any fungal genera, while MiCoP detected 3 of the 4 genera in the sample. The genus that MiCoP did not detect was not present in the NCBI RefSeq fungal database, so MiCoP did as well as possible given the state of the NCBI database. In terms of speed, MetaPhlAn2 processed the reads significantly faster than MiCoP, but MiCoP still processed the 4.9 million reads in a reasonable time of 2 hours and 55 minutes.

**Table 5.**
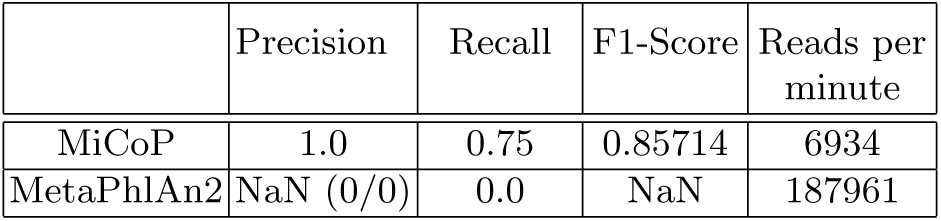
Comparison of the genus-level performance of MiCoP and MetaPhlAn2 on a mock fungal community consisting of 4 genera. MiCoP detects 3 of the 4 genera with no false positives, while MetaPhlAn2 detects nothing. Because MetaPhlAn2 has 0 true and false positives, precision cannot be computed. MetaPhlAn2 was faster, but MiCoP still finished in less than 3 hours.

|  | Precision | Recall | F1-Score | Reads per minute |
| --- | --- | --- | --- | --- |
| MiCoP | 1.0 | 0.75 | 0.85714 | 6934 |
| MetaPhlAn2 | NaN (0/0) | 0.0 | NaN | 187961 |

We then compared the performance of MetaPhlAn2 and MiCoP on the HMP data, using the parameter settings validated on the mock community datasets. Following an example provided by the MetaPhlAn authors, we downloaded 20 samples from the HMP, 10 from buccal mucosa and 10 from tongue dorsum. We analyzed each sample using MiCoP and MetaPhlAn2. Figures 2 to 5 show the results for MiCoP and MetaPhlAn2 for the relative abundances of fungi and viruses. Note that the relative abundances of fungi were computed with respect to the reads that could be identified as coming from fungi, and likewise for viruses, not with respect to the entire sample. MetaPhlAn2 estimates relative abundances for the entire sample; here, we recomputed the relative abundances for fungi and viruses relative to themselves. Results are shown in Figures 2, 3, 4, and 5.

**Fig. 2.**
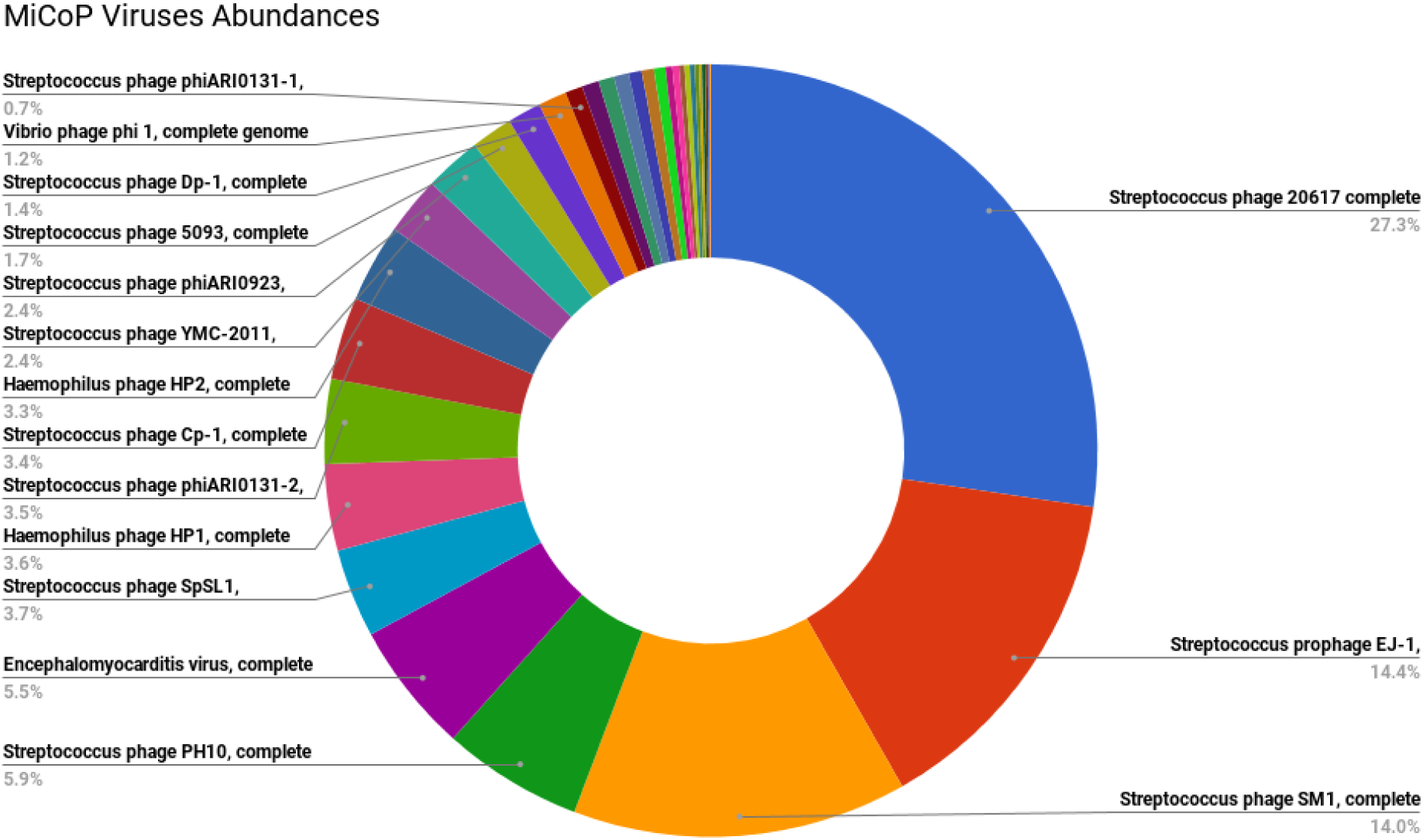
Abundance estimation for viruses when applying MiCoP to 20 Human Microbiome Project samples, 10 from buccal mucosa and 10 from tongue dorsum. MiCoP detects a total of 34 species present, with the sample being dominated by bacterial phages, particularly *Streptococcus* phages.

**Fig. 3.**
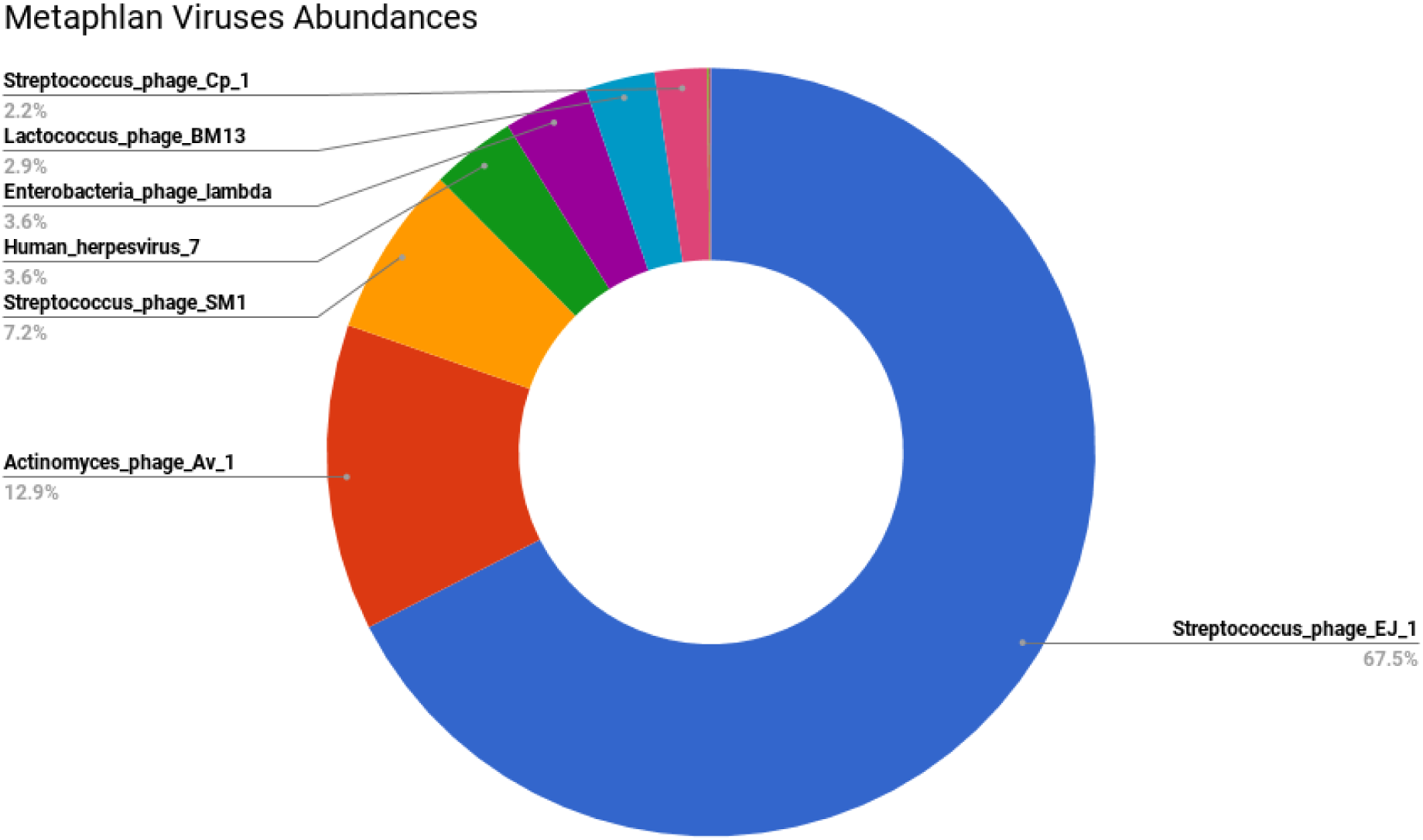
Abundance estimation for viruses when applying MetaPhlAn2 to 20 Human Microbiome Project samples, 10 from buccal mucosa and 10 from tongue dorsum. MetaPhlAn finds a much lower virome diversity than MiCoP, with only 12 species identified. The sample is again dominated by *Streptococcus* phages, but MetaPhlAn’s results suggest that there is only a single type of this phage dominating the sample, while MiCoP suggests that a wide variety of *Streptococcus* phages are present. MetaPhlAn’s results may stem from the reference bias issue explored in the simulation studies.

**Fig. 4.**
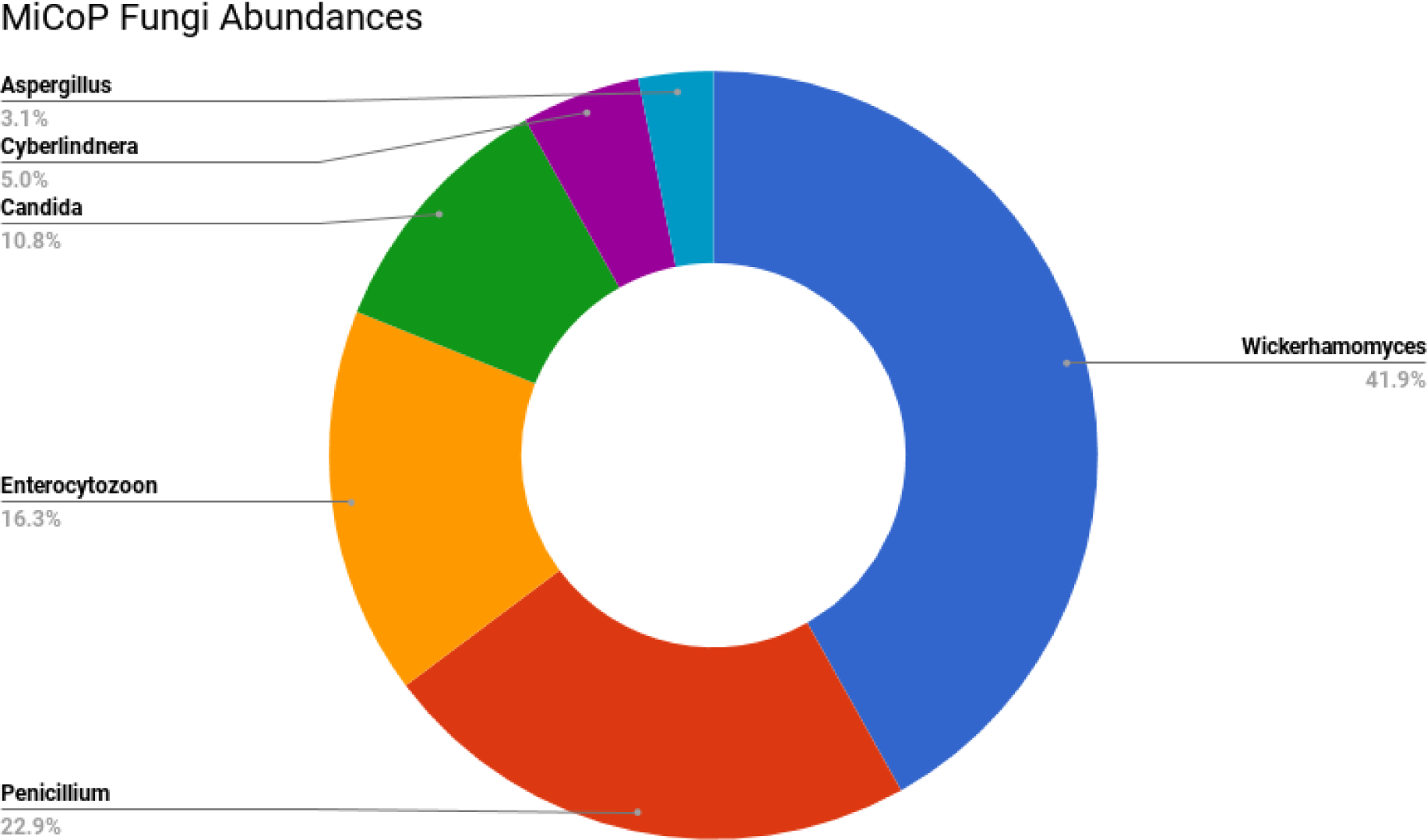
Abundance estimation for fungi when applying MiCoP to 2G Human Microbiome Project samples, 1G from buccal mucosa and 1G from tongue dorsum. MiCoP detects a total of 6 genera present.

**Fig. 5.**
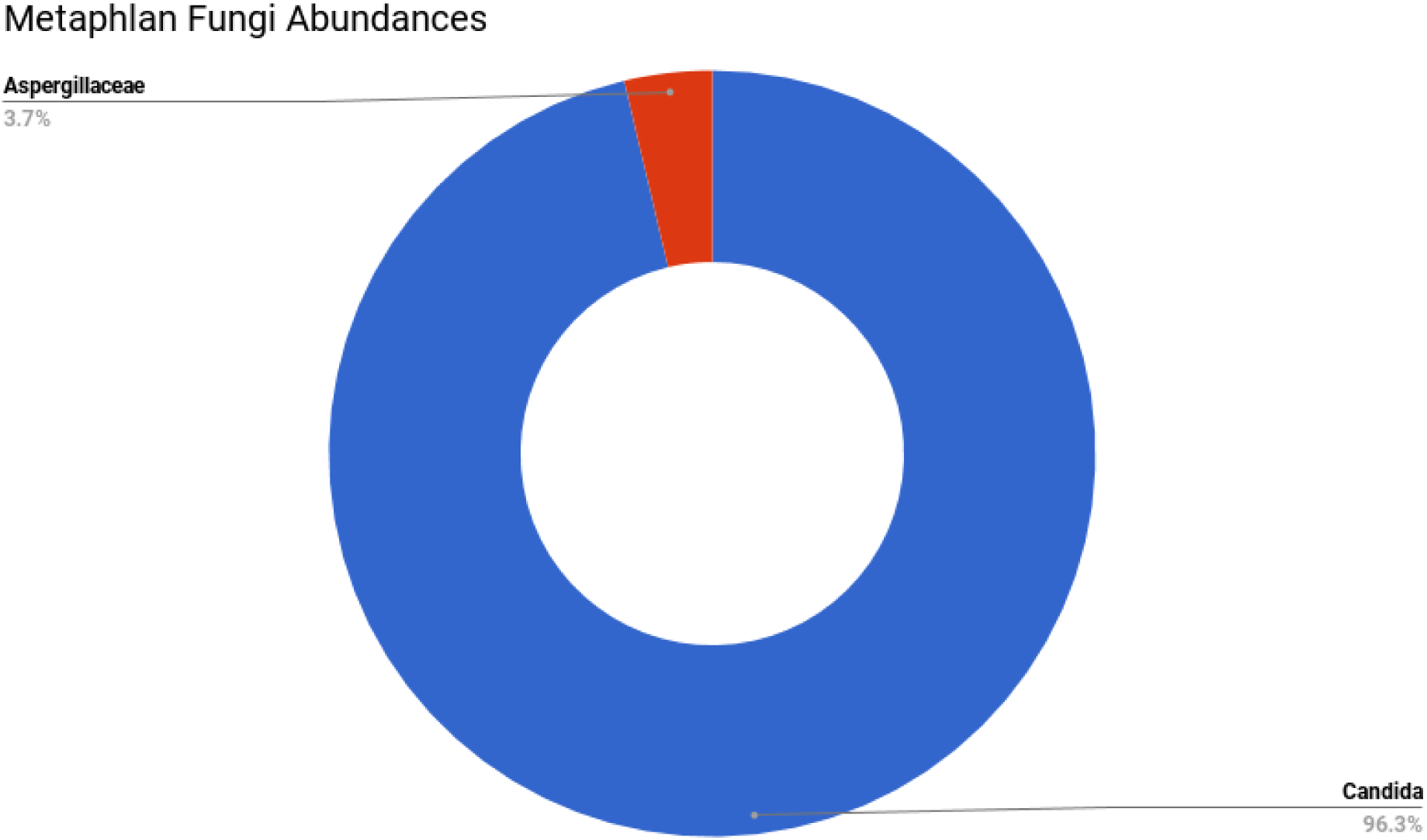
Abundance estimation for fungi when applying MetaPhlAn2 to 20 Human Microbiome Project samples, 10 from buccal mucosa and 10 from tongue dorsum. MetaPhlAn detects only two genera (*Candida* and *Aspergillaceae*), which are also present in MiCoP’s results. As the human oral eukaryome is known to be diverse [33–35], our results indicate that MiCoP captures the fungal community diversity better.

We found that MiCoP consistently identified a more diverse virome and eukaryome than MetaPhlAn across all samples and both body sites. For fungi, MetaPhlAn2 identified two genera in the HMP samples, *Candida* and *Aspergillaceae,* while MiCoP identified 6 genera, including the two identified by MetaPhlAn2. Additionally, while the *Candida* genus dominated the MetaPhlAn2 results with 96.3% abundance, genera identified by MiCoP were distributed in a more balanced manner. For viruses, MetaPhlAn identified 12 species present while MiCoP identified 34 species. Both results were dominated by *Streptococcus* phages, but MetaPhlAn’s results were dominated by *Streptococcus phage EJ1,* while MiCoP identified a diverse group of *Streptococcus* phages. As the human oral virome and eukaryome are known to be highly diverse [33–35], our results indicate that MiCoP is capturing more of the community diversity than MetaPhlAn. MiCoP can be used as an effective alternative to popular general-purpose metagenomic abundance estimation tools when a more comprehensive characterization of the human virome and eukaryome is desired.

## Discussion and Conclusion

MiCoP illustrates the benefits of a mapping-based approach for metagenomic analyses, especially of viral and eukaryotic species. Methods that are infeasible for the largest bacterial reference databases can be leveraged for smaller reference databases due to increased sensitivity. The mapping-based approach is a particular example of this, as it is more sensitive to viral and eukaryotic species and gives valuable coverage information that other methods do not provide. Generally speaking, we observe that different mapping methods tend to be optimized for different types of microbes, and many existing methods are less effective for non-bacterial species. We also note the issue of reference bias, which our simulations showed can significantly impact the performance of profiling methods. If users attempt to use existing methods with their default databases, they may not accurately detect non-bacterial species. Our real data analysis supports this view, as MiCoP identified more species than previous studies had reported. In terms of run time, MiCoP trades off speed for sensitivity, so Kraken and MetaPhlAn2 run faster. However, this difference is relatively minor (significantly less than an order of magnitude) when using viral reference databases such as the NCBI RefSeq viral genomes due to their small size. The difference is more pronounced with eukaryote reference databases, due to the large genome size of eukaryotes, and can be more than an order of magnitude. Thus, MiCoP is likely to scale better for viral data than for eukaryotes.

There are several potential future directions for MiCoP. One possible extension would be to add a precomputation method that reduces the reference database size by removing genomes that have no chance of being in a set of reads, using k-mer or MinHash based methods. This would enable MiCoP to run faster and use less memory, perhaps making it feasible to analyze large bacterial reference databases. Another possible direction involves assembly of sequences that were not mapped to any reference genome. This would allow for the detection of species that are not available in a reference database, but caution would have to be taken to avoid false discoveries. MiCoP promises to help researchers more comprehensively and accurately identify viral and eukaryotic species in metagenomic samples.

## Methods

### Reference database and mapping method

Any metagenomic profiling method that aims to classify sequence reads as belonging to certain reference genomes is dependent to a large extent on the reference database used [15]. We show empirical evidence of this reference bias in our results section. Choosing the reference database involves a tradeoff between smaller databases that result in lower sensitivity but can be searched faster, and larger databases that take longer to search but enable more accurate results. In general, increasingly powerful computer hardware and fast mapping algorithms have enabled searching of large reference databases in a reasonable amount of time [19,36]. Additionally, viral and eukaryotic reference databases are currently much smaller than bacterial reference databases, making mapping-based approaches feasible. We therefore performed analysis using the full NCBI RefSeq Viral and Fungal databases. In addition to database selection, the selection of the mapping algorithm used in a mapping-based approach heavily affects results. We evaluated several new and established mapping methods, including Megablast [21], BWA [23], Bowtie2 [37], and Diamond [38]. We found that BWA produced the best results overall, comparable in accuracy to Megablast but with a much faster run time that was feasible for large modern metagenomic sequencing datasets.

### Probabilistic assignment of multi-mapped reads

While classifying uniquely-mapped reads is trivial, proper assignment of multi-mapped reads has a major impact on results. BWA’s default setting randomly chooses which genome to assign multimapped reads to; this setting led to a large amount of false positives in our simulated datasets. However, simply discarding all multi-mapped reads leads to poor read utilization and negatively affects sensitivity and abundance estimation. Thus, a method for accurately assigning multi-mapped reads is of critical importance in read classification.

In the first stage of our two-stage approach, uniquely-mapped reads are classified according to the genome that they map to, and the relative read counts for each genome are then calculated. All multi-mapped reads, and the list of the genomes that they map to, are set aside during this stage. During the second stage, we assign multi-mapped reads to a genome with probability equal to the relative uniquely-mapped read counts for each of those genomes. A consequence of this approach is that genomes whose only mapped reads are multi-mapped will have no chance of reads being mapped to them, and reads that map only to species bearing no uniquely-mapped reads will not be mapped at all. We also filter out genomes with fewer than 10 uniquely-mapped reads, as there is insufficient evidence to indicate their presence; this heuristic technique has been successfully employed in previous studies [19]. In comparison to letting BWA randomly choose multi-mapped read classification, we observed that these filtering steps have a minor-to-negligible impact on the sensitivity and vastly increase precision on the species level by eliminating many false positives.

### Relative abundance estimation

Following the classification of reads to genomes, we estimate the relative abundances of each organism in the sample. Many read classification methods do not support this step, even though indicating the actual pervasiveness of different species in a sample is more informative than pure read counting. To do this, we normalize the read counts for each genome by the length of the genome. We then normalize the adjusted counts of each genome by the sum of the adjusted counts, so that all species abundances sum up to 100%.

### Simulated datasets

Simulated reads from viral and eukaryotic genomes were generated using Grinder [39]. We used two different settings of simulated microbial communities, low complexity communities and high complexity communities. The parameters for these communities were set in accordance with the simulations performed by the CAMI consortium benchmark [15]. Low complexity communities consisted of 40 genomes with abundances selected from a lognormal distribution with mean 1 and standard deviation 2, then normalized such that they total 100%. High complexity communities were similarly produced, except with 544 genomes, mean 1.5, and standard deviation 1. All viral and eukaryotic genome simulations consisted of 1 million reads with lengths picked from a normal distribution with mean 150 and standard deviation 15.

## List of Abbreviations

MiCoP: Microbial Community Profiling
BWA: Burrows-Wheeler Aligner
NCBI: National Center for Biotechnology Information
RefSeq: NCBI Reference Sequence Database
MetaPhlAn: Metagenomic Phylogenetic Analysis
HMP: Human Microbiome Project
CAMI: Critical Assessment of Metagenome Interpretation
TP/FP/FN: True Positives / False Positives / False Negatives

## Declarations

### Ethics approval and consent to participate

Not applicable

## Consent for publication

Not applicable

## Availability of data and material

The code, data, and documentation for this study is publicly available on GitHub at: https://github.com/smangul1/MiCoP

BWA and the NCBI RefSeq databases are also publicly available online via their respective websites.

## Authors’ contributions

NL wrote the manuscript and code and carried out the experiments. SM developed the project and experiments with NL. MA, IM, NW, DK, and EE collaborated on the development of the project and its direction and goals. NW provided interpretation of the viral results.

## Acknowledgements

The authors would like to thank Lana Martin for her editorial assistance with the manuscript.

